# Nutrient Challenges in a Changing Atmosphere: Investigating Biomass Growth and Mineral Concentration Changes in Soybean Plants under Elevated CO_2_

**DOI:** 10.1101/2024.08.02.606357

**Authors:** Ravneet Kaur, Mary Durstock, Stephen A. Prior, G. Brett Runion, Elizabeth A. Ainsworth, Ivan Baxter, Alvaro Sanz-Sáez, Courtney P. Leisner

## Abstract

Rising atmospheric CO_2_ levels, projected to reach ∼650 ppm by 2050, threaten the nutritional value of food crops. This rise is expected to increase biomass yield in C_3_ plants through enhanced photosynthesis and water-use efficiency. However, elevated CO_2_ (eCO2) reduces protein, nitrogen, and essential minerals like zinc (Zn) and iron (Fe) in plant leaves and seeds, posing a global nutrition risk. We conducted an experiment using Open Top Chambers to examine the response of three soybean cultivars (Clark, Flyer, and Loda) to ambient (∼410 ppm) and eCO_2_ (∼610 ppm) conditions. These cultivars were selected due to their contrasting responses to eCO_2_. Measurements of physiological parameters (i.e., biomass, and nutrient concentration) were taken at different growth stages. Our results showed that eCO_2_ increased carbon assimilation, leading to higher aboveground biomass and seed yield (through increased seed number) while root biomass remained unchanged. eCO_2_ also reduced stomatal conductance and transpiration. There was a significant decrease in seed nutrient concentration at maturity, particularly iron (Fe), phosphorous (P), potassium (K), and magnesium (Mg), in plants grown in eCO_2_. These findings suggest that increased yield, reduced transpiration, and unchanged root biomass are key drivers of nutrient dilution in seeds under eCO_2_.

## INTRODUCTION

Since the industrial revolution in the late 1800s anthropogenic emissions of carbon dioxide (CO_2_) have increased (Friedlingstein *et al*., 2023; IPCC, 2023). Consequently, the concentration of CO_2_ in the Earth’s atmosphere has reached unprecedented levels on a global scale (Friedlingstein *et al*., 2023). There are several studies providing evidence of increased leaf area index, biomass and yield because of increasing atmospheric CO_2_ concentrations (eCO_2_) in C_3_ plants (Ferris *et al*., 1999; Dermody *et al*., 2006; Digrado *et al*., 2024). This increase in biomass is caused by the enhanced rate of photosynthesis with simultaneous decrease in stomatal conductance which drives the increase in water-use efficiency (WUE) in C_3_ plants, resulting in a "fertilization" effect (Drake *et al*., 1997; Long *et al*., 2006; Loladze, 2014; Myers *et al*., 2014; Ainsworth & Long, 2021). Consequently, the projected rise in eCO_2_ to ∼ 650 ppm by 2050 has the potential to positively impact world food production and address the needs of a growing population (Ciais *et al*., 2014).

Several studies have shown that the increased plant growth due to eCO_2_ is accompanied by a significant reduction in protein, nitrogen and several other mineral nutrients in plant leaves and seeds (Högy & Fangmeier, 2009; Loladze, 2014; Dietterich *et al*., 2015; Myers *et al*., 2017; Soares *et al*., 2019; Ainsworth & Long, 2021). Previous studies have shown that C_3_ grains and legumes, when cultivated in future eCO_2_ conditions projected for 2050, have reduced zinc (Zn) and iron (Fe) levels (Loladze, 2014; Myers *et al*., 2014). The reported decrease in mineral nutrition is of consequence, as much of the world population relies on C_3_ grains and legumes for their primary Zn and Fe intake (Tulchinsky, 2010; Myers *et al*., 2014). This decrease in micronutrients in C_3_ grains and legumes has the potential therefore to increase essential micronutrient deficiencies in developing and developed nations, impacting around 2 billion individuals (Tulchinsky, 2010).

This change in micronutrients due to eCO_2_ is associated with “hidden hunger”, which is defined as plant-based diets that meet caloric, but not nutritional needs (Kennedy, 2002; Welch & Graham, 2005). Previous work has shown that plants grown in eCO_2_ are adding a teaspoon of total nonstructural carbohydrates (TNC) (about 5g of a starch-and-sugar mixture) per 100g of dry plant mass (Loladze, 2014). The effect on TNC:protein and TNC:mineral ratios is similar to the stoichiometric impact of adding a spoonful of carbohydrates to every 100g of dry plant tissue (Loladze, 2014). To combat this “hidden hunger” (Kennedy, 2002; Welch & Graham, 2005) a greater understanding of the physiological mechanisms responsible for reduced mineral concentration in seeds and other plant organs under higher levels of eCO_2_ is needed.

There are several hypothesized mechanisms associated with the reduction in nutrient concentration in C_3_ plants grown in eCO_2_. This includes 1) a reduction in transpiration in leaves and thus reduced bulk flow of nutrients (Mcgrath & Lobell, 2013); 2) an increase in the carbohydrate and fiber content, leading to a mineral dilution in seeds and other plant parts (Poorter *et al*., 1997; Gifford *et al*., 2000; Taub & Wang, 2008; Taub *et al*., 2008; Chaturvedi *et al*., 2017); 3) a reduction in root mineral absorption due to changes in the structure of roots (Beidler *et al*., 2015) and an alteration in the expression of mineral transporters in root tissue (Taub & Wang, 2008; Leakey *et al*., 2009; Jauregui *et al*., 2016), and 4); it has been demonstrated that conditions reducing photorespiration, like high eCO_2_, also restrict nitrate uptake and assimilation (Searles & Bloom, 2003; Rachmilevitch *et al*., 2004; Bloom *et al*., 2010). It is important however, to note that there is a lack of empirical data supporting all these hypotheses and that these hypothesized mechanisms are not mutually exclusive with one another. Further work is needed to experimentally examine these hypotheses to determine the most likely mechanisms contributing to reduced nutrient concentration.

Soybean (*Glycine max* L. Merr.) is an extensively cultivated crop and is a model legume species that has notable phenotypic variation related to biomass accumulation and yield responses to eCO_2_ (Bishop *et al*., 2015; Sanz Sáez *et al*., 2017). Furthermore, soybean cultivars that exhibited an increased yield response to eCO_2_ levels also exhibited a notable decrease in crucial mineral nutrients, specifically Fe and Zn, in their seeds (Loladze, 2014; Myers *et al*., 2014; Bishop *et al*., 2015; Aranjuelo *et al*., 2015). The mechanisms associated with this phenotypic variation in yield and mineral nutrient response to eCO_2_ in soybeans are largely unknown, however (Schmutz *et al*., 2010; Myers *et al*., 2014; Parvin *et al*., 2019; Soares *et al*., 2019). To begin investigating these unknowns, an experiment was conducted on the effects of elevated CO_2_ levels on soybean physiology and plant tissue mineral concentration. Results from this work will help identify the physiological factors responsible for nutrient loss under high CO_2_ conditions. Additionally, these findings can be used to help guide future molecular biology experiments to test the transcriptomic responses associated with these physiological changes playing an important role in developing molecular tools for creating more climate-resilient crops.

## MATERIALS AND METHODS

### Plant Material and Experimental Conditions

Soybeans were grown at the USDA-ARS National Soil Dynamics Laboratory in Auburn, AL, in open-top chambers (OTC). OTC consisted of a cylindrical, aluminum metal frame that was 3 m wide and 2.4 m tall, with the bottom half covered with clear plastic, allowing the sunlight to penetrate and reach the plants (Rogers *et al*., 1983; Runion *et al*., 2008). The double-walled plastic chamber cover consisted of 2.5 cm of perforations in the inner plastic wall, allowing gas distribution into the chamber. Two atmospheric eCO_2_ were used across eight chambers, four at ambient (∼410 ppm) and four elevated (ambient + 200 ppm CO_2_) during daylight hours. Three soybean cultivars (Clark, Flyer, and Loda) were chosen based on their different yield and/or nutrient accumulation responses under eCO_2_: Loda had high strong yield response; Flyer showed decreased Zn accumulation in seeds; and Clark showed no change in Zn in seeds (Myers *et al*., 2014; Sanz Sáez *et al*., 2017; Soba *et al*., 2020; Digrado *et al*., 2024). Chambers were set up in a randomized complete block design (*n* = 4), and each chamber contained four plants from each of the three cultivars.

Seeds were inoculated with commercial *Bradyrhizobium japonicum* (N-dure, Verdesian Inc., Cary, NC, https://vlsci.com/products/n-dure/) and germinated in a greenhouse on May 6, 2021. On May 10, 2021, seedlings were transplanted into 20-liter black containers filled with soil from the E.V. Smith Research Station (Shorter, AL). The soil is classified as sandy loam, consisting of 23.6% silt and clay, 76.4% sand, 3.2% clay, and 20.4% silt with a pH of 6.1. Immediately following transplanting, containers were placed in the OTC. Following soil test recommendations from Auburn University Soil Testing Laboratory, 1 gram of potash was applied to each container and 2 grams of Miracle Gro to ensure sufficient nutrient availability. Plants were watered daily with a drip tape irrigation system that applied 1.9 liters of water every other day for the first four weeks and every day afterwards to avoid drought stress.

### Leaf Gas Exchange, Crop Growth, and Harvest Measurements

Leaf CO_2_ assimilation (*A*) and stomatal conductance (*g_s_*) were measured using an infrared gas analyzer (LI-6800, LI-COR Biosciences, Lincoln, NE). Measurements were conducted during midday hours (10:00 a.m.–2:00 p.m.) on the most recently fully expanded leaf located at the top of the canopy. These measurements were taken 34 days after planting (DAP) on June 9 (V5 vegetative stage) and 91 DAP (full seed, R6 developmental stage) (Fehr *et al*., 1971) on August 5. To ensure accuracy, the light intensity and temperature inside the leaf cuvette were adjusted to match the ambient conditions. The LI-6800 was used to measure the ambient light intensity. The relative humidity within the leaf cuvette was maintained at 60–70% and the concentration of CO_2_ inside the cuvette was set to match that of the ambient or elevated OTC conditions. Gas exchange measurements were averaged from two plants per cultivar per OTC per time point.

Biomass sampling occurred 70 DAP (pod filling, R5 developmental stage) on July 15^th^ and 126 DAP (maturity, R8 developmental stage) on September 9th. At the pod-filling stage (70 DAP) seed count were measured immediately at harvest. Both aboveground biomass (shoots, leaves, and pods) and belowground biomass (roots) were collected at 70 DAP. The samples were oven-dried for at least 72 hours at 60 °C and then weighed. At maturity (126 DAP), aboveground biomass (stems and pods) was collected. These samples were also dried for 72 h at 60 °C and then weighed. Biomass measurements were averaged from two plants per cultivar per OTC per time point. Harvest index (HI) (HI=seed yield/ (seed yield + aboveground biomass) was also calculated for both 70 and 126 DAP harvest.

Nutrient analysis was also completed at 70 and 126 DAP. Aboveground (leaf, stem, and pod) and belowground (root) samples were analyzed at 70 DAP, and aboveground (stem and pods) samples were analyzed at 126 DAP. Dried tissue samples were weighted and ground. The samples were sent to Waters Agricultural Laboratory, Inc. in Camilla, GA, for nutrient concentration analysis. Macronutrient concentrations (%) of nitrogen (N), phosphorus (P), potassium (K), magnesium (Mg), sulfur (S), calcium (Ca), and micronutrient concentrations (ppm) of boron (B), iron (Fe), zinc (Zn), manganese (Mn) and copper (Cu) were determined using inductively coupled plasma mass spectroscopy (ICP-MS). Samples were averaged from two plants per cultivar per OTC at both time points. Nutrient concentration of aboveground biomass samples was calculated as the sum across leaf, stem and pod tissues. Nutrient uptake for macronutrients (g/total biomass per tissue per plant) and micronutrients (mg/total biomass per tissue per plant) was calculated from nutrient concentration and biomass data. The percent change in measurements for elevated vs ambient CO_2_ was calculated as ((elevated-ambient)/elevated) *100.

### Statistical Analysis of physiological data and data visualization

Statistical analysis of the physiological data (gas exchange, biomass, and nutrient analysis) was conducted using a mixed model procedure of SAS (Littell *et al*., 1996). The cultivars and CO_2_ treatments were considered fixed effects, while blocks were considered random effects. When the fixed effect of cultivar, CO_2_ or their interaction was significant, Tukey’s honest significant differences post-hoc test was performed to compare means (LSMEANS, SAS 9.4, SAS Institute, Cary, NC, USA). Plots were made using ggplot2 in R (v.3.4.4; Wickham *et al*. 2016), and Microsoft^®^ Excel.

## RESULTS

### Gas exchange parameters were impacted by the eCO_2_ treatment

At 34 DAP, midday stomatal conductance (*g_s_*) (mol H_2_O m^-2^ s^-1^) significantly decreased in plants grown in eCO_2_ when averaged across all cultivars (Fig. 1A). Midday stomatal conductance (*g_s_*) decreased by 76.3% in Clark, 82.4% in Flyer, and 73.5% in Loda but main effect of cultivar was not significant (Fig. 1A). At 34 DAP midday carbon assimilation (*A*) (µmol CO_2_ m^-2^ s^-1^) significantly increased in plants grown in eCO_2_ when averaged across all cultivars but main effect of cultivar was not significant (Fig. 1B). Specifically, *A* increased by 18.6% in Clark, 9.4% in Flyer, and 15% in Loda (Fig. 1B). A similar trend was observed at 91 DAP as *g_s_* was significantly decreased under eCO_2_ when averaged across all cultivars (Fig. 1C) with both eCO_2_ and cultivar as significant main effects. *g_s_* was decreased by 15.4%, 31%, and 15% in Clark, Flyer and Loda, respectively, under eCO_2_ (Fig. 1C). At 91 DAP, *A* significantly increased in plants grown in eCO_2_ when averaged across all cultivars (Fig. 1D). *A* was increased by 6.4%, 18.1% and 19 % in Clark, Flyer and Loda, respectively and had significant cultivar main effect (Fig. 1D). There was also a trend for a CO_2_ by cultivar interaction (*p = 0.*07) at 91 DAP; Flyer and Loda had a significant increase in photosynthesis under eCO_2_, while Clark did not (Fig. 1D).

**Figure 1.**
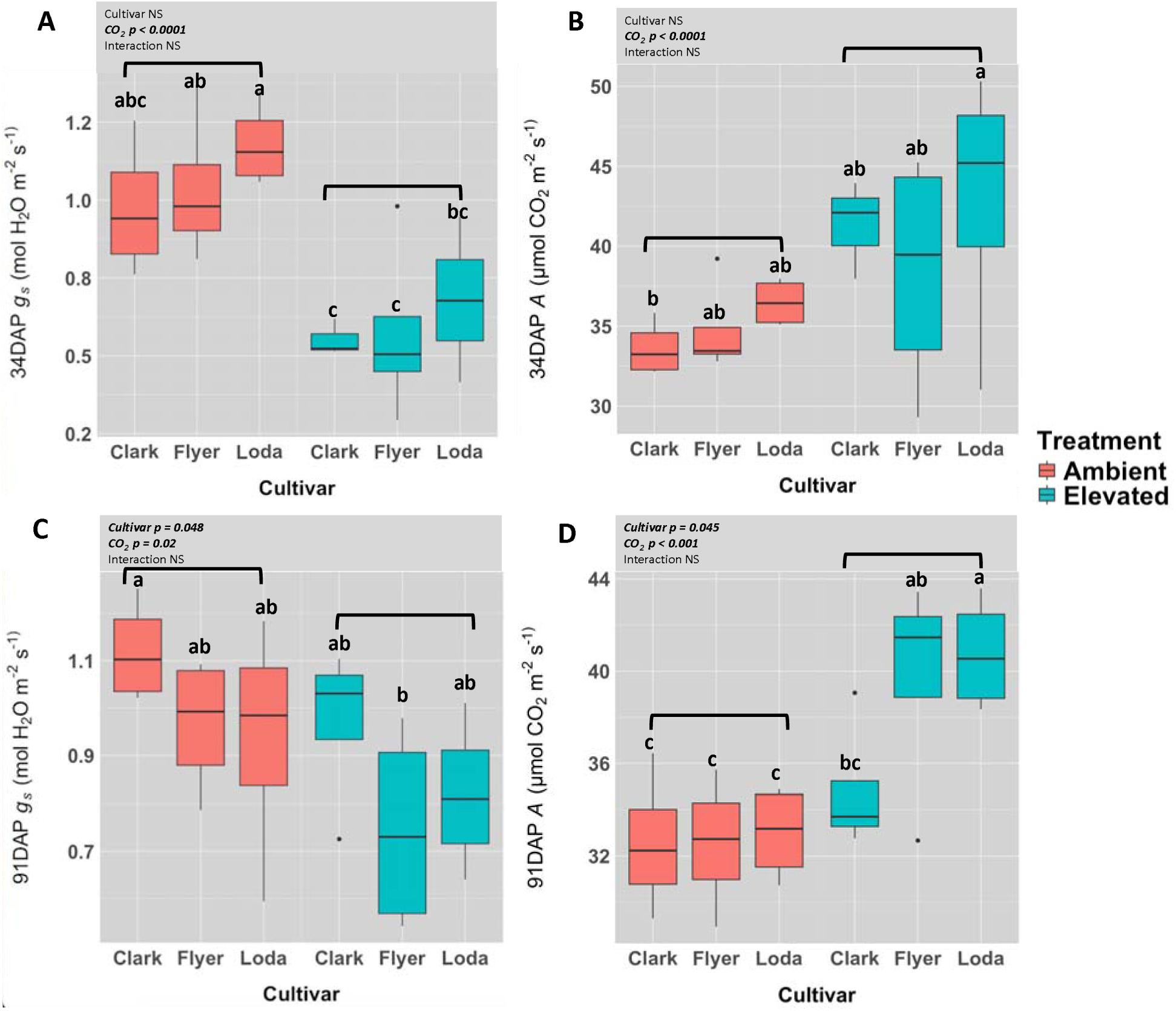
The effect of eCO_2_ on (A) stomatal conductance *(g_s_)* and (B) carbon assimilation *(A)* rate during 34 DAP and (C) stomatal conductance *(g_s_)* and (D) carbon assimilation rate *(A)* 91 DAP time points. Different letters indicate significant differences across treatments. Brackets represent significance (*p* < 0.05) of main effect of CO_2_ from the two-way ANOVA and are provided for clarity. NS represents no significance. Results of main effect statistics are placed in the upper corners.

### Biomass of aboveground but not belowground plant tissues increased under eCO_2_ conditions

At 70 DAP aboveground biomass was significantly increased under eCO_2_ levels when averaged across all cultivars, but there was no significant interaction between cultivar and CO_2_ (Fig. 2A, Table S1a). Aboveground biomass increased by 19.43%, 20.28% and 22.46% in Clark, Flyer and Loda, respectively. When averaged across all cultivars leaf biomass was also significantly increased under eCO_2_ (Fig. 3A). There was also a significant cultivar main effect for leaf biomass with Loda having lower biomass than Clark and Flyer (Fig. 3A). At 70 DAP, root biomass was affected only by cultivar; Loda had significantly lower root biomass than both Clark and Flyer (Fig. 3B Table S1a).

**Figure 2.**
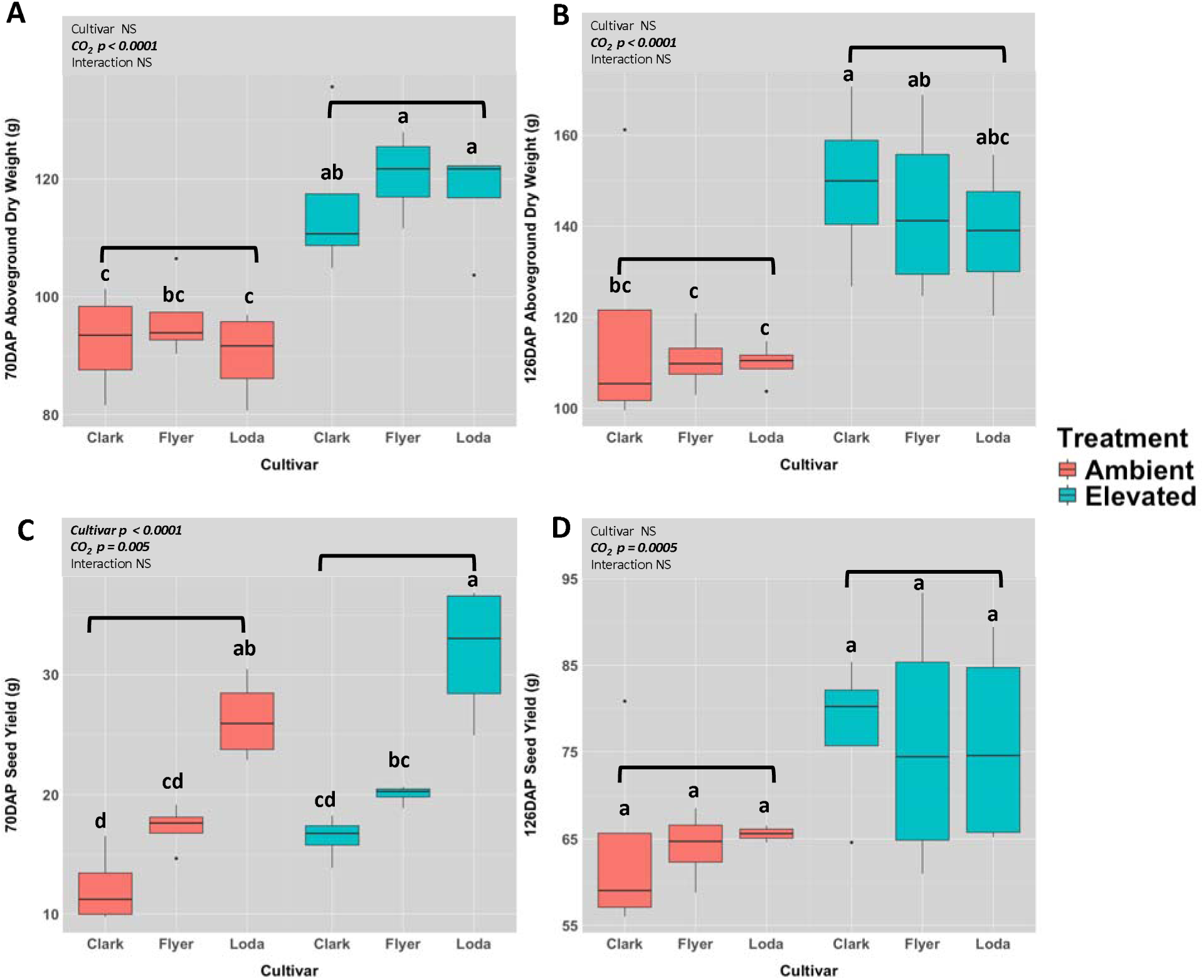
The effect of eCO_2_ on aboveground biomass and total seed yield. at (A) 70 DAP (B) 126 DAP and on seed yield at (C) 70 DAP and (D) 126 DAP. Different letters indicate significant differences across treatments. Brackets represent significance (*p* < 0.05) of main effect of CO_2_ from the two-way ANOVA and are provided for clarity. Results of main effect statistics are placed in the upper corners. NS represents no significance.

**Figure 3.**
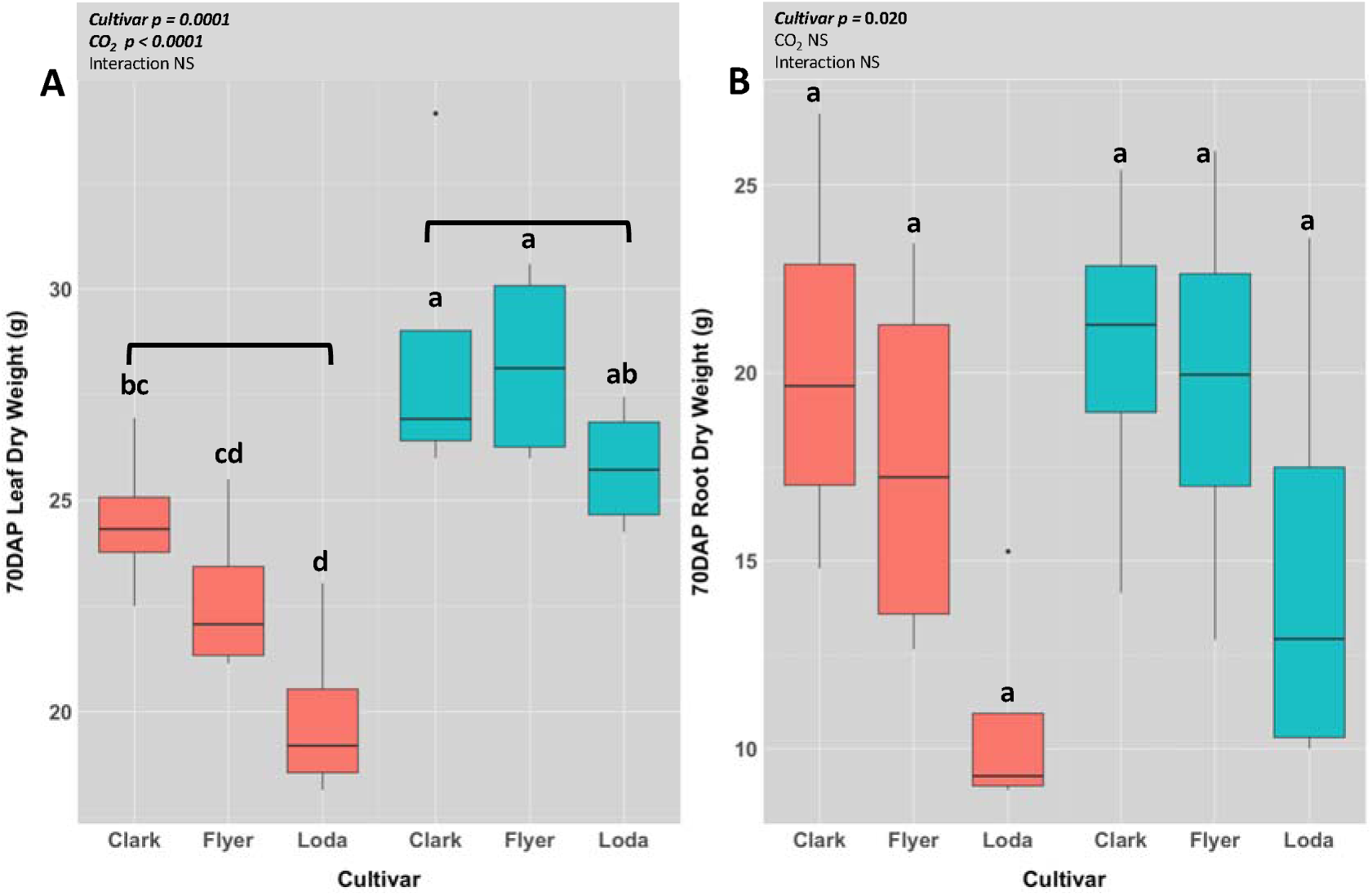
The effect of eCO_2_ on (A) leaf dry weight and (B) root dry weight at 70 DAP. Leaf biomass per plant of each cultivar at 70 DAP in ambient and eCO_2_. Different letters indicate significant differences across treatments. Brackets represent significance (*p* < 0.05) of main effect of CO_2_ from the two-way ANOVA and are provided for clarity. Results of main effect statistics are placed in the upper corners. NS represents no significance.

At 70 DAP seed yield increased under eCO_2_ when averaged across cultivars (Fig. 2C, Table S1a). There was also a significant cultivar effect on seed yield; Loda had significantly higher seed yield than Clark and Flyer, regardless of treatment (Fig. 2C). Per seed dry weight was not affected by elevated CO_2_ and varied significantly only with cultivar at 70 DAP (Fig. S1A). Per seed dry weight was significantly higher for Loda than other cultivars (Fig. S1A). At 70 DAP total seed number was higher under eCO_2_ and was also significantly affected by cultivar (Fig. S2A). Total seed number was significantly higher in Flyer than Clark, regardless of CO_2_ treatment (Fig. S2A). There was no effect of CO_2_ on HI at 70 DAP; however, a significant cultivar main effect was present (Fig. S3A). Loda had significantly higher HI than other cultivars regardless of CO_2_ treatments.

At 126 DAP, aboveground biomass (pod and stem) was significantly increased under eCO_2_ when averaged across cultivars (Fig. 2B, Table S1b). Flyer (21.5%) and Clark (22.39%) showed a significant increase and Loda (19.72%) showed a marginally significant increase in aboveground biomass under eCO_2_ compared to ambient CO_2_ (Fig. 2B, Table S1b). Seed yield had a 14.46% increase in plants grown in eCO_2_ when averaged across cultivars (Fig. 2D, Table S1b). Per seed dry weight only varied significantly with cultivar (Fig. S1B). Clark and Loda had significantly higher per seed dry weight than Flyer regardless of CO_2_ level (Fig. S1B). Total seed number significantly increased in plants grown in eCO_2_ when averaged across all cultivars (Fig. S2B). There was also a significant main effect of cultivar on total seed number at 126 DAP, with Flyer having higher seed numbers than the other cultivars (Fig. S2B). There was significant reduction in HI under eCO_2_ with CO_2_ and cultivar as significant main effect (Fig. S3B).

### Nutrient concentration in different plant tissues changed under eCO_2_ conditions

#### Macronutrients

The concentration of macronutrients (N, P, K, Mg, Ca, and S) was assessed at the 70 DAP and 126 DAP. At 70 DAP nutrient concentration was measured in root, seed, and aboveground biomass (combination of leaf, pod and stem). At 126 DAP, macronutrient concentration was measured in stems and seeds only. At 70 DAP the concentration of macronutrients in seeds varied significantly only with cultivar, except for S which had both cultivar and CO_2_ as significant main effects (Fig. 4A; Table S2). Additionally, N concentration had a significant CO_2_ x cultivar interaction (Fig. 4A). N had a decrease of 1.1% in Clark and 2.4% in Flyer, and an increase of 1.5% in Loda in eCO_2_. Leaf macronutrient concentration did not change with eCO_2_ or cultivar (Table S2). In roots no macronutrient concentration showed a significant change with eCO_2_ except for a significant increase in Mg (5.7-13.4%), a significant decrease in S (5-21%) and a marginally significant decrease in K (*p* = 0.07; 7.2-38%) when averaged across cultivars (Table S2). In aboveground tissues N, Mg and Ca varied significantly with cultivar. K was significantly reduced with eCO_2_ while P and S varied significantly by both eCO_2_ and cultivar (Table S2). Nutrient uptake of macronutrients increased in leaves (6.3%-38.43%) under eCO_2_ at 70 DAP when averaged across all cultivars and was also significantly impacted by cultivar regardless of CO_2_ treatment (Fig. S4A; Table S4). The leaves of cultivar Loda showed significantly higher uptake of Mg, Ca, and S, whereas Flyer had a significant increase in Ca and S uptake under eCO_2_ (Fig. S4A; Table S4). At 70 DAP nutrient uptake of only N and Mg were significantly increased by eCO_2_ when averaged across cultivars in roots (Fig. S4B; Table S4), while Ca (*p = 0.052*) showed a marginally significant increase in nutrient uptake in eCO_2_ averaged across all cultivars (Fig. S4B; Table S4). Uptake of all macronutrients in roots varied significantly with cultivar at 70DAP except for Cu (*p* = 0.081) (Fig. S4B; Table S4).

**Figure 4.**
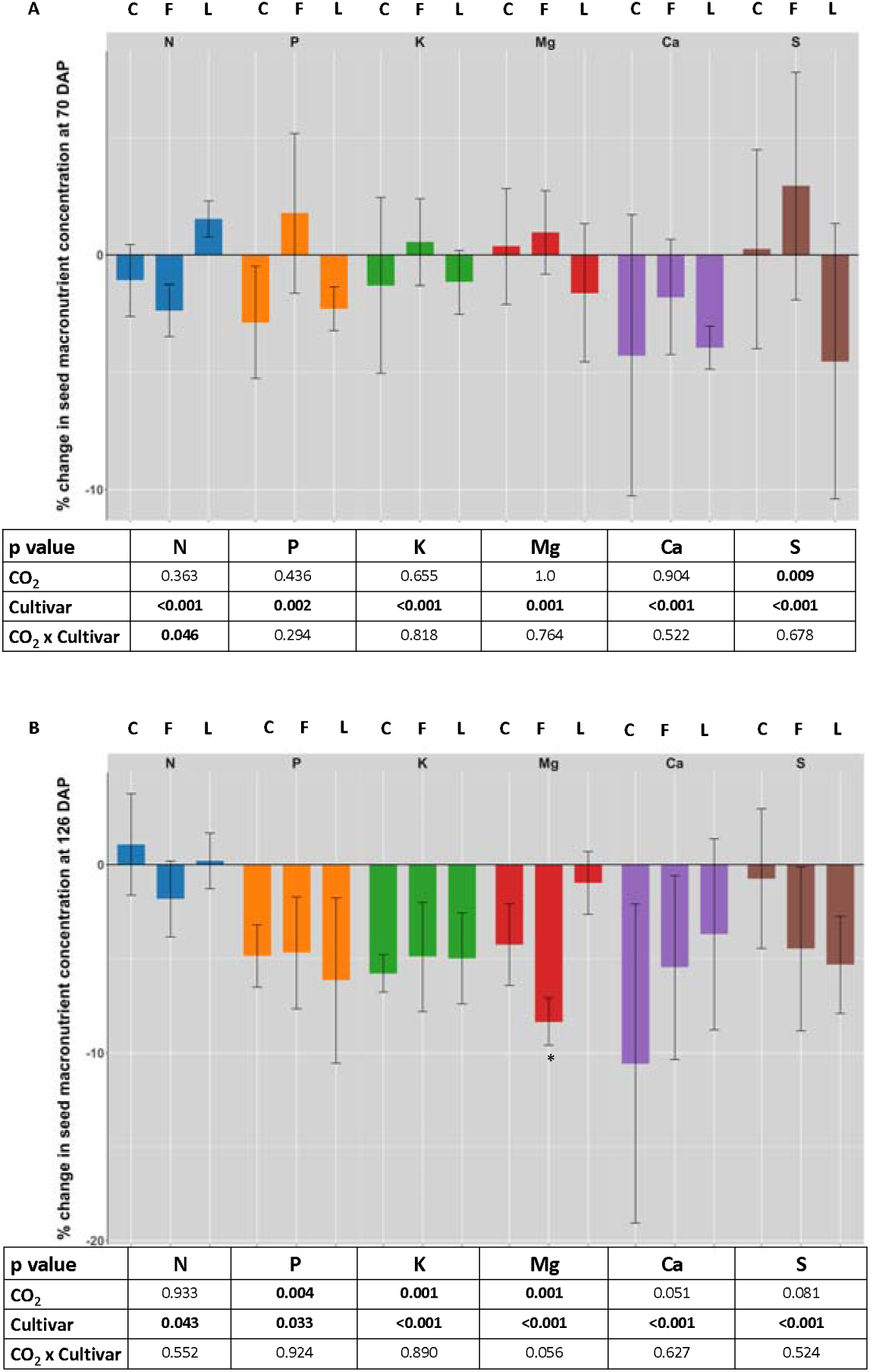
Percent change (%) at eCO_2_ versus ambient CO_2_ of seed macronutrient concentration in Clark (C), Flyer (F) and Loda (L). (A) The percent change in macronutrient (N, P, K, Mg, Ca, and S) concentration in seed, during (A) 70 DAP and (B) 126 DAP. Percent change was calculated as ((elevated-ambient)/elevated) *100. Asterisks indicate significant differences (*p* < 0.05) between elevated and ambient measured/absolute values (for values see Table S2-3). The *p* values are from two-way ANOVA of the absolute values. Results of main effect statistics are placed in the table below the figure.

At 126 DAP all seed macronutrients showed a significant main effect of cultivar (Fig. 4B; Table S3). There was significant decrease caused by eCO_2_ for P, K, and Mg when averaged across all cultivars (Fig 4B). Reduction in P, K, and Mg concentrations in seed under eCO_2_ varied from 0.7% - 10.6% across cultivars. There was a moderately significant reduction in Ca (*p* = 0.051) in plants grown in eCO_2_ when averaged across cultivars (Fig. 4B). There was also a moderately significant CO_2_ x cultivar interaction for Mg (*p* = 0.056) (Fig. 4B; Table S3). Mg decreased in Flyer significantly with eCO_2_ (Fig. 4B). In aboveground tissue macronutrients such as P, Ca and S varied significantly by cultivar (Table S3). None of the macronutrients in aboveground biomass showed significant change with eCO_2_ (Table S3). At 126 DAP seeds showed an increase (4.1% - 154.2%) for uptake of all macronutrients under eCO_2_, with Cu having a moderately significant increase (*p* = 0056). Mg and Ca also varied significantly with cultivar as significant main effect (Fig. 6; Table S4).

#### Micronutrients

The concentration of micronutrients (B, Zn, Mn, Fe, and Cu) was also measured 70 and 126 DAP. At 70 DAP nutrient concentration was measured in root, seed, and aboveground biomass and at 126 DAP nutrient concentration was measured in stems and seeds only. All seed micronutrients at 70 DAP showed a significant main effect of cultivar, and Zn, B, and Cu were significantly decreased by eCO_2_ when averaged across cultivars (Fig. 5A; Table S2). Mn had a slight increase in eCO_2_ when averaged across cultivars (*p* = 0.077). There was also a significant CO_2_ x cultivar interaction for Zn and Cu in seeds at 70 DAP (Fig. 5A). The concentration of Zn in Clark was reduced by 15.7% in eCO_2,_ and Cu in Loda was reduced by 14.8% under eCO_2_. The concentration of B was significantly reduced in Clark under eCO_2_ by 23.1% (Fig. 5A; Table S2). At 70 DAP B, Zn and Cu were the only micronutrients which varied in aboveground tissue with eCO_2_ as significant main effect (Table S2). None of the micronutrients in leaf tissue changed significantly with eCO_2_ or cultivar (Table S2). Concentration of Zn, Fe and Mn in roots varied with cultivar as significant main effect whereas concentration of B and Cu did not show significant change with eCO_2_ or cultivar (Table S2).

**Figure 5.**
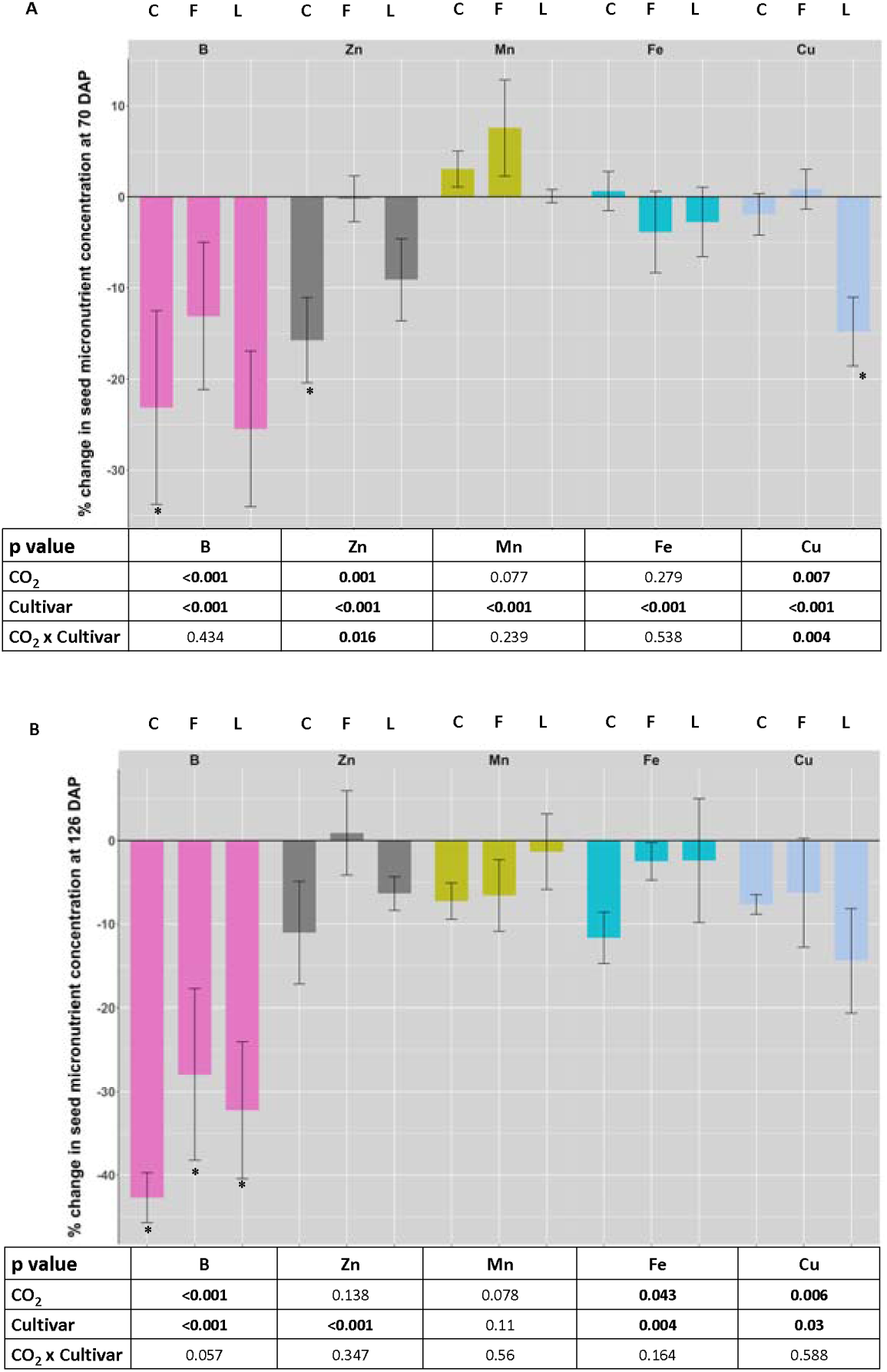
Percent change (%) at eCO_2_ versus ambient CO_2_ of the seed micronutrient concentration in Clark (C), Flyer (F) and Loda (L). The percent change in micronutrient (Zn, Fe, B, Mn, and Cu) concentration in seed at (A) 70 DAP and (B) 126 DAP. Percent change was calculated as ((elevated-ambient)/elevated) *100. Asterisks indicate significant differences (*p* < 0.05) from the two-way ANOVA between elevated and ambient measured/absolute values (see Table S2-3).

Nutrient uptake for micronutrients was also measured at 70 DAP. Nutrient uptake of micronutrients was significantly increased by eCO_2_ in leaves at 70 DAP, with the except of Zn and Fe (Fig. S4A; Table 4). Only B and Cu had a significant cultivar main effect, with Mn having a moderate cultivar main effect (*p* = 0.065) (Fig. S4A; Table S4). B showed a significant increase in Loda and Cu showed a significant increase in Flyer under eCO_2_ (Fig. S4A; Table S4). The micronutrient uptake in root tissue at 70 DAP varied with cultivar as significant main effect except Fe (*p* = 0.055) and Cu (*p* = 0.081) which had marginally significant effect of cultivar (Fig. S4B; Table S4). There was no significant impact of eCO_2_ on nutrient uptake of micronutrients in roots at 70 DAP (Fig. S4B; Table S4).

At 126 DAP, seed concentrations of Fe, B, and Cu significantly decreased under eCO_2_ when averaged across cultivars (Fig. 5B; Table S3). Additionally, there was a significant effect of cultivar on Zn, Fe, B, and Cu levels. Both Fe and Cu concentrations were reduced across all cultivars, with Fe decreasing by 11.7 in Clark, 2.5% in Flyer, and 2.4% in Loda, and Cu decreasing by 7.7% in Clark, 6.3% in Flyer, and 14.4% in Loda. Zn concentrations were reduced only in Clark (11.04%) and Loda (6.35%) under eCO_2_. B also showed a marginally significant CO_2_ x cultivar interaction (*p* = 0.057), with B concentrations significantly reduced in Clark (42%), Flyer (27%), and Loda (32%) under eCO_2_. Although there was no significant effect of CO_2_ or cultivar on Mn concentration, there was a general decrease in Mn concentration with eCO_2_ when averaged across cultivars (*p* = 0.078). B was the only micronutrient that decreased in aboveground tissue with both eCO_2_ and cultivar as significant main effects (Fig. 6; Table S3. Uptake of all micronutrients except B increased in seeds at 126 DAP under eCO_2_ (Fig. 6). Reduction in B uptake was significantly impacted by both CO_2_ and cultivar and B was decreased in eCO_2_. Finally, Zn uptake was significantly impacted by cultivar only (Fig. 6; Table S3).

**Figure 6.**
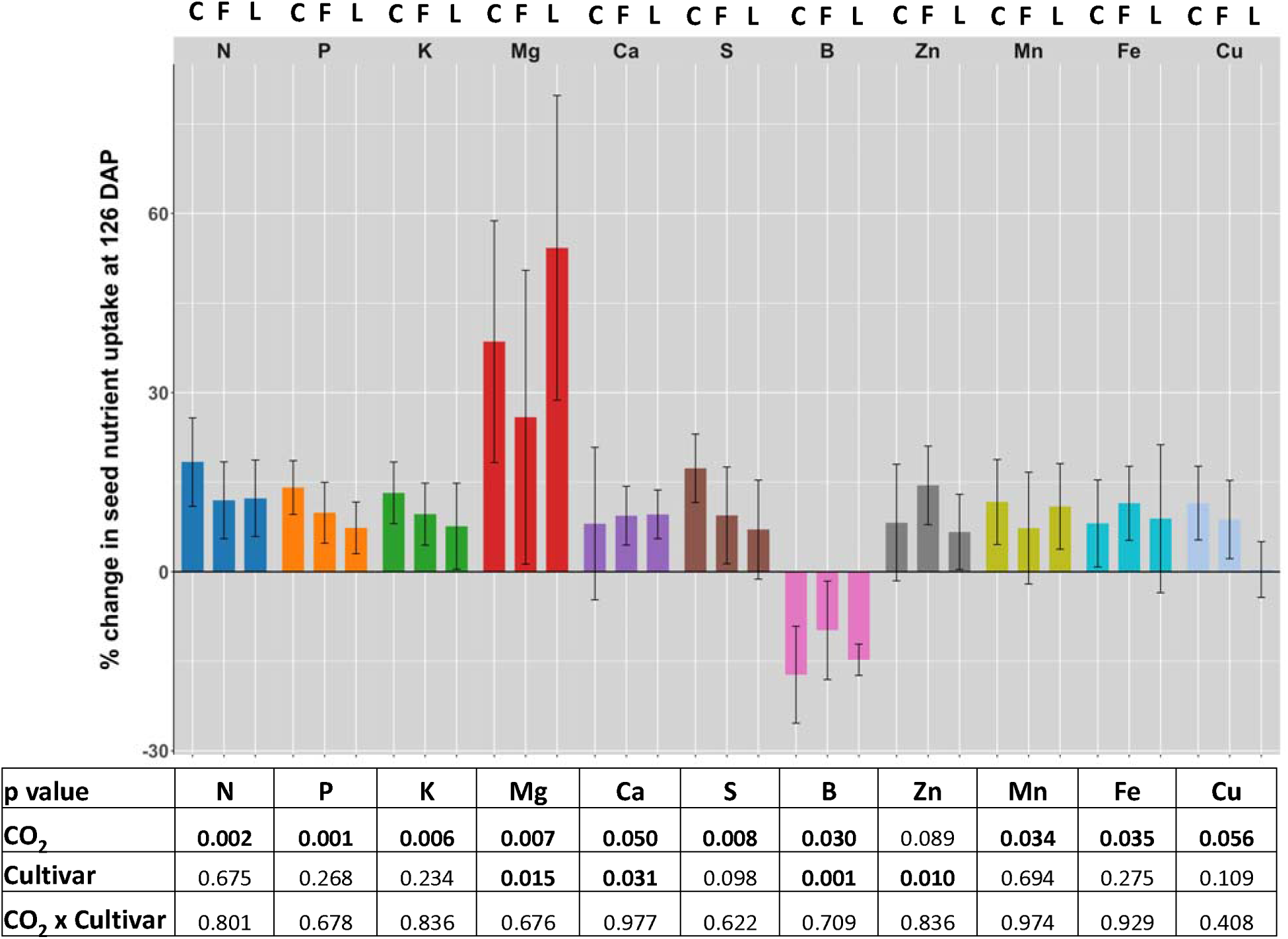
Percent change (%) at eCO_2_ versus ambient CO_2_ of the seed nutrient uptake in Clark (C), Flyer (F) and Loda (L). The percent change in macro (g/total biomass per tissue per plant) and micro (mg/total biomass per tissue per plant) nutrient uptake (macronutrients: N, P, K, Mg, Ca, S; micronutrients: B, Zn, Mn, Fe and Cu) in seed at 126 DAP. Percent change was calculated as ((elevated-ambient)/elevated) *100. Asterisks indicate significant differences between elevated and ambient measured/absolute values (see Table S4). The *p* values are from two-way ANOVA of the absolute values.

## DISCUSSION

The aim of this study was to explore the physiological mechanisms underlying plant nutrient responses to eCO_2_ levels. We selected soybean, a key C_3_ commodity crop and a model for legume species, due to its significant role in ecosystem services through atmospheric nitrogen fixation via symbiotic relationships with microorganisms (Schmutz *et al*., 2010). Our research focused on three soybean cultivars—Clark, Flyer, and Loda—chosen for their contrasting phenotypes under eCO_2_ conditions (Myers *et al*., 2014; Sanz Sáez *et al*., 2017). Flyer and Loda were known to show reduction in Zn concentration (Myers *et al*., 2014). Loda was previously documented to show increased yield and Clark was documented to be non-responsive under eCO_2_ (Bishop *et al*., 2015). This approach allowed us to investigate how different cultivars respond to increased CO_2_ and the resulting impacts on nutrient uptake and assimilation.

### Increased carbon assimilation and WUE under eCO_2_ resulted in increased yield due to increase in seed number

The eCO2 treatments significantly enhanced *A* and reduced *g_s_*in soybean plants, leading to improved WUE (Fig. 1). At both 34 and 91 DAP, *A* increased notably across all cultivars, with Flyer and Loda showing the most significant enhancements under eCO_2_ conditions (Fig. 1B-1D). This increase in *A* was accompanied by a substantial reduction in *g_s_*, indicating improved WUE (Fig. 1A-1C). The increased photosynthetic efficiency under eCO_2_ conditions contributed to greater biomass accumulation in aboveground plant tissues (Fig. 2A) at 70, as well as at 126 DAP (Fig. 2B).

Furthermore, eCO_2_ led to a significant increase in total seed yield, primarily driven by an increase in seed number rather than individual seed weight (Fig. 2C-D; Fig. S1-S2). This suggests that the enhanced yield observed under eCO_2_ can be attributed to the increased number of seeds produced per plant, highlighting the role of improved carbon assimilation and WUE in driving yield increases in soybean. The findings underscore the potential of eCO_2_ to enhance crop productivity through physiological adaptations that boost photosynthesis and optimize water usage, resulting in more seeds and total yield per plant (Fig. 1; Fig. 2B). Our results align with previously reported studies showing increased *A* and WUE resulting in stimulated increased biomass and yield in soybean (Ferris *et al*., 1999). Previous work also found increased seed yield under eCO_2_ was attributable to increased pod number and total seed number per plant, causing an increase in total seed yield per plant (Rogers *et al*., 1984; Li *et al*., 2013).

### Reduced seed nutrient concentration was observed when plants were grown in eCO_2_

The concentrations of macronutrients like P, K, and Mg, as well as micronutrients such as Fe, B, and Cu were significantly decreased in seeds at maturity (126 DAP) in soybean plants grown in eCO_2_ (Fig. 4-5 and Table S2-S3). There was also a moderately significant reduction in Ca and Mn concentrations in seeds under eCO_2_ at maturity (Fig. 4-5). Mg is essential for the functioning of ribulose 1,5-bisphosphate carboxylase/oxygenase (Rubisco) and chlorophyll (McGrath & Lobell 2013), and the increased carbon assimilation associated with eCO_2_ levels during the 70 DAP could potentially explain the reduced Mg concentration in seeds. Previous studies have also observed a similar reduction in Cu concentration under eCO_2_, while Mg has been previously shown to increase in eCO_2_ in fresh edible soybean varieties (Li *et al*., 2018). Taken together, findings from this study suggest that the observed increase in seed yield, driven by the higher number of seeds, might contribute to the observed mineral dilution across the total seeds produced by each plant. Similar results have been seen previously where increased yield either due to eCO_2_ or breeding has resulted in reduced P, K, Mg, Fe, B, Zn, and Cu concentration (Monasterio & Graham 2000; Mcgrath & Lobell 2013; Myers *et al*., 2014).

### Cultivar-specific responses to eCO_2_ were observed

eCO□ significantly increased seed yield in Loda, primarily due to an increased number of seeds rather than individual seed weight (Fig. S3-S4). This increase in yield was associated with a substantial rise in *A* and a notable reduction in *g_s_*, leading to improved WUE (Fig. 1A and 1C). Despite the higher yield, Loda exhibited a consistent reduction in both macro- and micronutrient concentrations at 70 and 126 DAP. Previous studies have demonstrated a strong yield increase for Loda under eCO□ (Bishop *et al*., 2015; Sanz Sáez *et al*., 2017; Digrado *et al*., 2024). This trend was evident across multiple growth stages, highlighting the challenge of nutrient dilution in high-yielding cultivars under future atmospheric conditions.

Clark showed a negative percent change in nutrient concentration in seeds at 126 DAP for macronutrients like P, K, Mg, S, and Ca, and micronutrients such as Zn, Fe, B, Mn, and Cu. It also showed increased seed yield with eCO□ (Fig. 2D; Fig. 4-5; Table S2, S3). Flyer exhibited an increase in aboveground biomass at both 70 and 126 DAP (Fig. 2A and 2B), along with a notable reduction in key nutrients such as P, K, and Mg in seeds at 126 DAP (Fig. 4-5). However, we observed almost no change in Zn concentration at 70 DAP and a slight positive percent change in Zn concentration in Flyer at 126 DAP (Fig. 4-5).

Previously, Clark has been shown to be non-responsive to eCO□ in terms of yield and seed nutrient concentration, and Flyer has been known to show a significant reduction in Zn concentration (Myers *et al*., 2014; Bishop *et al*., 2015). A study by Bishop et al., (2015) examining the impact of eCO_2_ on nine cultivars of soybean (including Clark) reported a significant impact of genotype by year interaction on seed yield, however. Previous studies have also shown the complexity of yield and nutrient concentration responses in soybean cultivars grown under different environmental stresses, geographical locations, and during different years (Köhler *et al*., 2019; Digrado *et al*., 2024). For example, a recent study found that when soybean plants were grown in eCO_2_ and elevated temperature there no observed yield gains or nutrient losses (Köhler *et al*., 2019). This indicates that the observed cultivar-specific responses observed in our study may differ from previous studies due to genotype x environmental interactions. This is an important consideration for future work investigating cultivar-specific impacts of eCO_2_ on nutrient concentration in soybean.

### Lack of response of root biomass with eCO_2_ may be contributing to decreases in nutrient concentration in seeds

Our study reports an increase in seed yield and aboveground biomass at both 70 DAP and 126 DAP (Fig. 4 and Table S1). Leaf biomass at 70 DAP increased with eCO_2_ in all genotypes (Fig. 3A). Leaf biomass and nutrient uptake of all nutrients except Zn and Fe at 70 DAP increased with eCO_2_ in all genotypes (Fig. 3A; Fig. S4A). Although the effect of the cultivar on nutrient uptake by leaves was significant for most nutrients, and marginally significant for Mn (*p* = 0.065) it was not significant for Zn and Fe. Under eCO_2_, the uptake of all nutrients, except for Zn and Fe, increased significantly when averaged across all cultivars (Fig. S4A; Table S4). However, root biomass measured at 70 DAP (Fig. 3B) did not show any change under eCO_2_. Additionally, we observed increases in the uptake of all macro- and micronutrients in seeds (except for a decrease in B) and leaves (except Zn and Fe) under eCO_2_ (Fig. 6; Fig. S4A), In roots however, but, there was no significant response in nutrient uptake, except for increase N, Mg, and Ca (*p* = 0.052) to eCO_2_ (Fig. S4B).

Root architecture, morphology, and physiology play a primary role in water and nutrient acquisition from the soil (Wang *et al*., 2006) and could therefore play an important role in nutrient concentration in seeds. Previous work has found that eCO_2_ increased root biomass in non-nodulating soybeans (Rogers *et al*., 1992), while Van Vuuren et al., (1997) found eCO_2_ delayed root development in spring wheat. We hypothesize that if eCO_2_ results in delayed root development during growth stages associated with pod filling and nutrient accumulation in seeds (Bender *et al*., 2015), it could lead to reduced nutrient absorption and transport by roots. It is of note that while our study was done in pots, we used 20-liter pots which have been shown to be of sufficient size to not impact root biomass in soybean (Ainsworth *et al*., 2002; Poorter *et al*., 2012). Future work focusing on soybean root biomass changes under eCO_2_ in field conditions is needed to better understand how root biomass changes under eCO_2_ and the impact this may have on nutrient accumulation in seeds.

## CONCLUSION

Overall, findings of this study enhance our understanding of how eCO_2_ impacts the nutrient concentration in soybean seeds. By measuring physiological parameters such as carbon assimilation (*A*) and stomatal conductance (*g_s_*), and the biomass of individual tissues like leaf, seed, stem, and root we highlighted the role of *g_s_*and nutrient uptake and transport, particularly in roots. Our work supports the claim that eCO_2_ leads to a reduction in micronutrient concentrations in seeds and there is significant cultivar variation in this response. Furthermore, we observed likely genotype x-environment interactions in cultivar-specific responses to eCO_2_, namely in the change in Zn and other nutrients in Clark which was previously hypothesized to be non-responsive to eCO_2_. We propose that an increased number of seeds, rather than increased per seed weight, contributes to the reduction in seed nutrient concentration due to mineral dilution. Additionally, eCO_2_ resulted in increased *A* and increased biomass of leaves, stems, and seeds but did not impact root biomass. The root nutrient uptake of only N, Mg and Ca increased significantly under eCO_2_. This suggests that roots could not compensate for the increase in *A* by enhancing nutrient absorption, uptake and transport, leading to reduced nutrient concentration in seeds. Future research should aim to enhance our understanding of how nutrient transport and absorption by roots are affected under elevated CO_2_ conditions.

## ACKNOWLEDGEMENTS

We would like to thank Collin Modelski for assisting in various activities of data collection. This work is supported by a USDA NIFA award (#2022-67013-36126) to CPL.

## CONFLICT OF INTEREST STATEMENT

The authors declare no conflicts of interest.

## DATA AVAILABILITY STATEMENT

Data from this research will be made available upon request.

## Supplemental Figures

**Figure S1.**
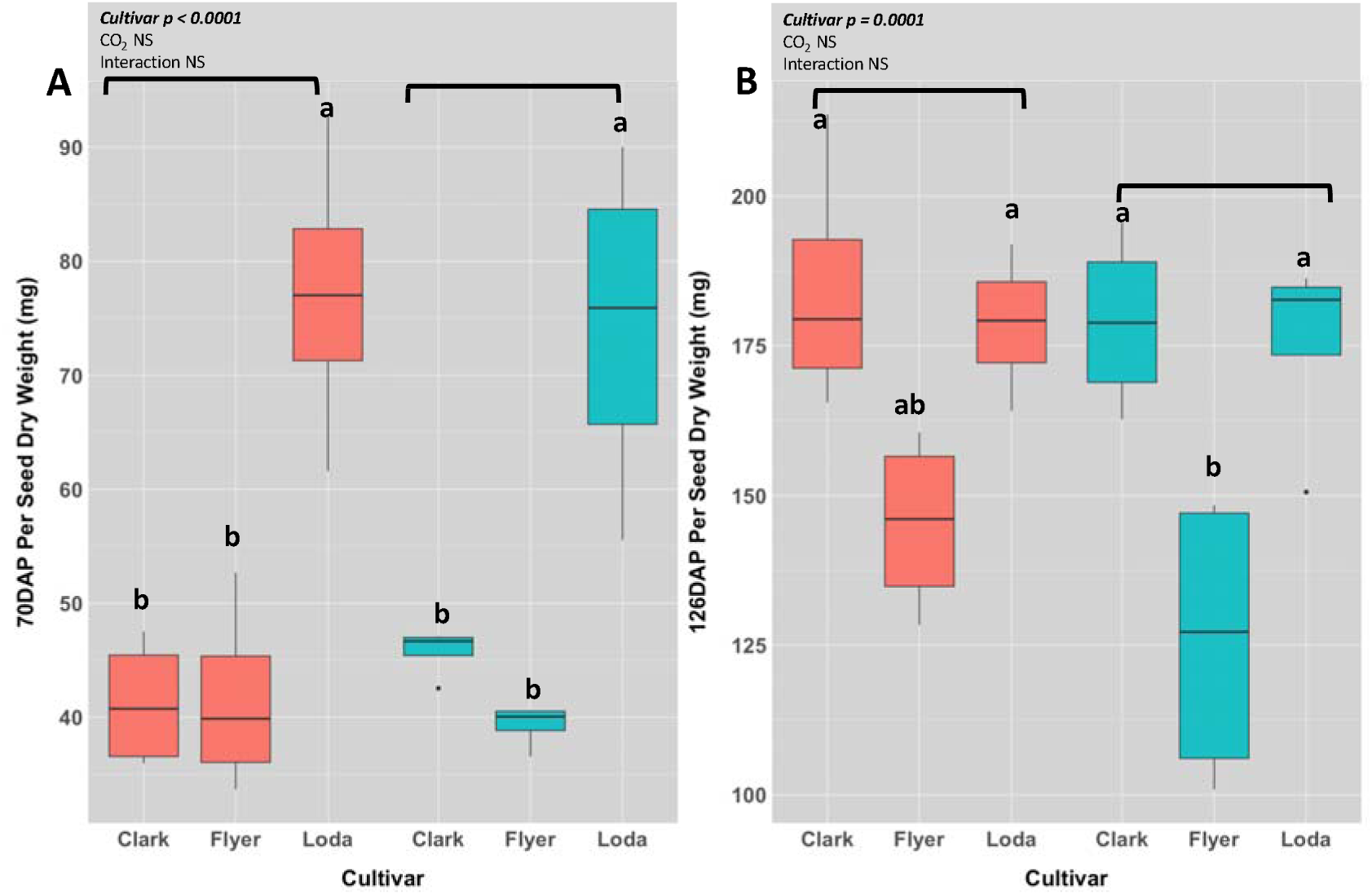
The effect of eCO_2_ on per seed dry weight (A) 70 DAP (B) 126 DAP. Per seed dry weight of each cultivar at 70 DAP in ambient and eCO_2_. Different letters indicate significant differences across treatments. Brackets represent significance (*p* < 0.05) of main effect of CO_2_ from the two-way ANOVA and are provided for clarity. Results of main effect statistics are placed in the upper corners. NS represents no significance.

**Figure S2.**
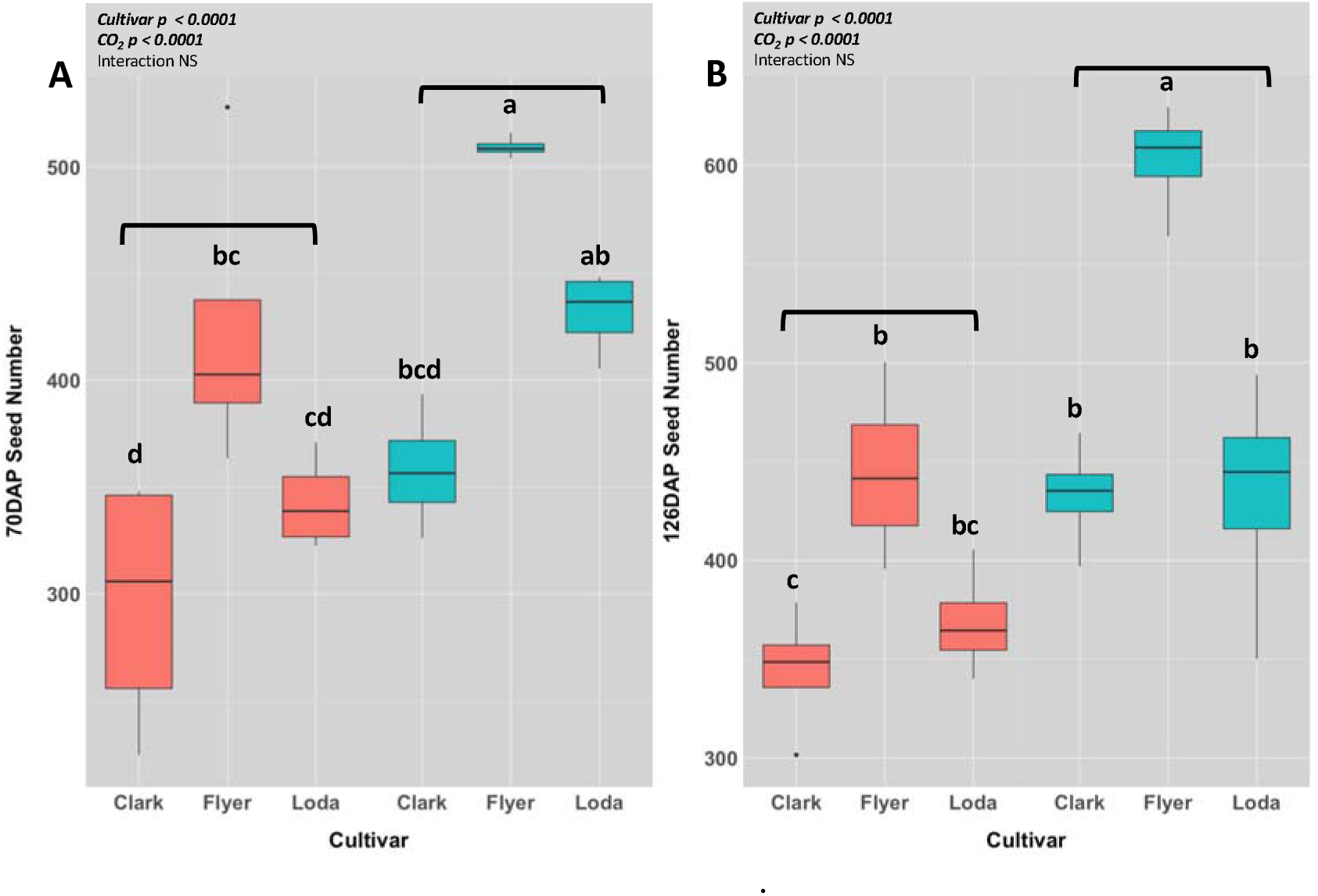
The effect of eCO_2_ on seed number at (A) 70 DAP and (B) 126 DAP. Seed number per plant of each cultivar at 70 DAP in ambient and eCO_2_. Different letters indicate significant differences across treatments. Brackets represent significance (*p* < 0.05) of main effect of CO_2_ from the two-way ANOVA and are provided for clarity. Results of main effect statistics are placed in the upper corners. NS represents no significance.

**Figure S3.**
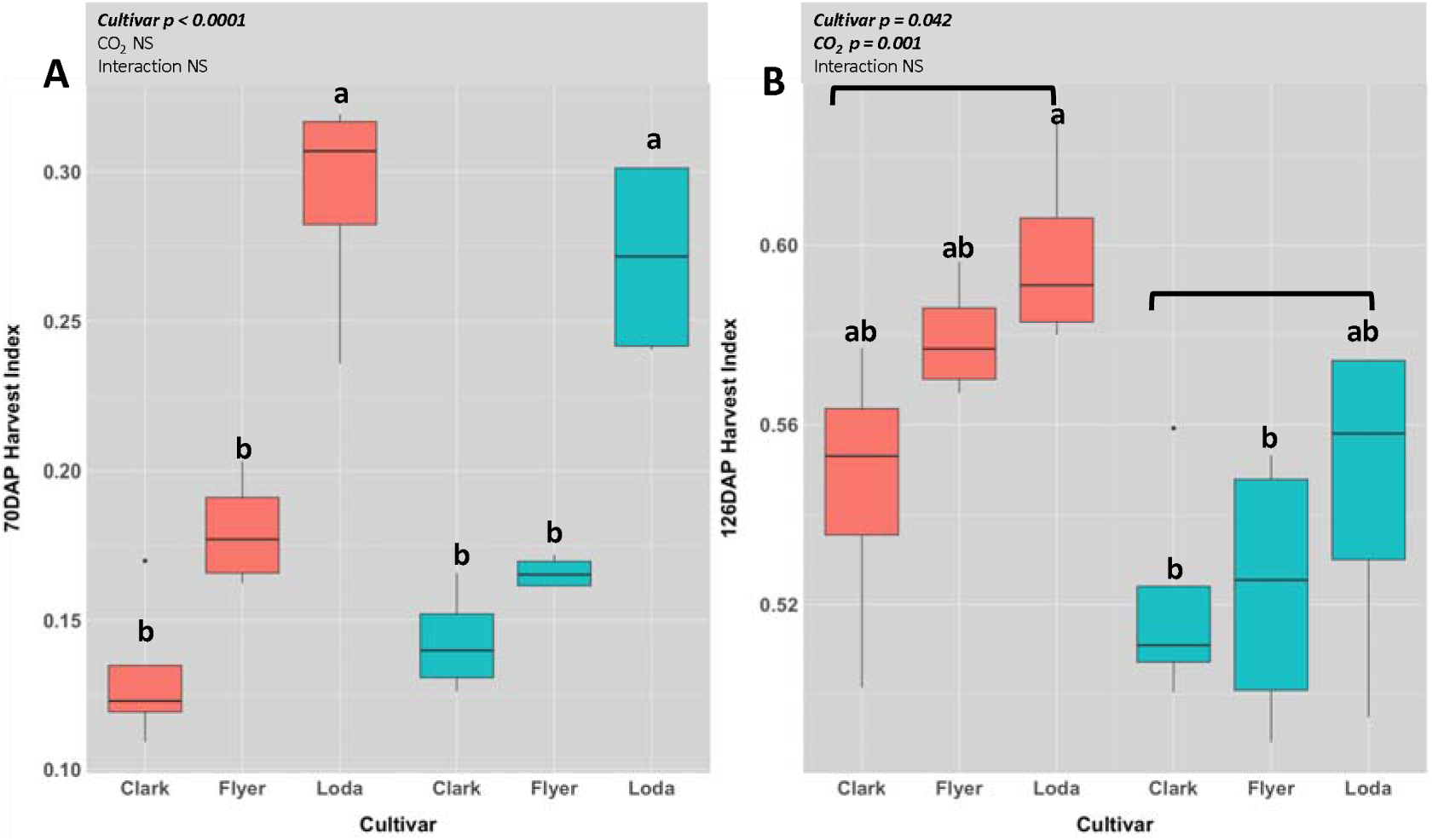
The effect of eCO_2_ on harvest index at (A) 70 DAP and (B) 126 DAP. Harvest index per plant of each cultivar at 70 DAP in ambient and eCO_2_. Different letters indicate significant differences across treatments. Brackets represent significance (*p* < 0.05) of main effect of CO_2_ from the two-way ANOVA and are provided for clarity. Results of main effect statistics are placed in the upper corners. NS represents no significance.

**Figure S4.**
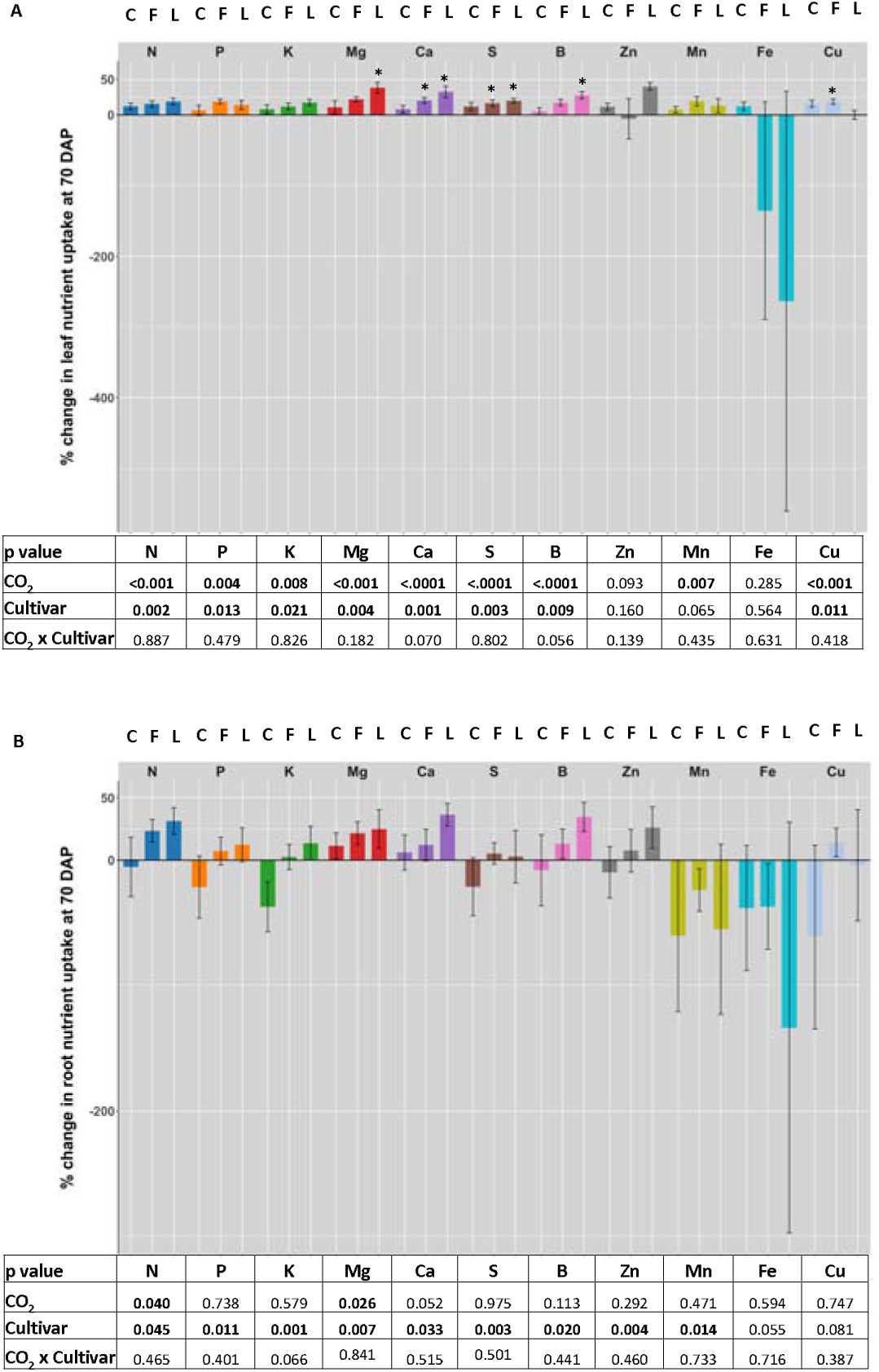
Percent change (%) at eCO_2_ versus ambient CO_2_ of the root and leaf nutrient uptake in Clark (C), Flyer (F) and Loda (L). The percent change in macro-(g/total biomass per tissue per plant) and micronutrient (mg/total biomass per tissue per plant) uptake (macronutrients: N, P, K, Mg, Ca, S; micronutrients: B, Zn, Mn, Fe and Cu) in (A) leaf and (B) root tissue at 70 DAP. Percent change was calculated as ((elevated-ambient)/elevated) *100. Asterisks indicate significant differences between elevated and ambient measured/absolute values (see Table S4). The *p* values are from two-way ANOVA of the absolute values.

## Notes

### Competing Interest Statement

The authors have declared no competing interest.

